# Plasma Aβ ratios in autosomal dominant Alzheimer’s disease: the influence of genotype

**DOI:** 10.1101/2021.02.11.430756

**Authors:** Antoinette O’Connor, Josef Pannee, Teresa Poole, Charles Arber, Erik Portelius, Imogen Swift, Amanda J Heslegrave, Emily Abel, Nanet Willumsen, Helen Rice, Philip SJ Weston, Natalie S Ryan, James M Polke, Jennifer M Nicholas, Chris Frost, Simon Mead, Selina Wray, Lucía Chávez-Gutiérrez, Kaj Blennow, Henrik Zetterberg, Nick C Fox

**Affiliations:** Dementia Research Centre, UCL Queen Square Institute of Neurology, London, United Kingdom; UK Dementia Research Institute at UCL, London, United Kingdom; Department of Psychiatry and Neurochemistry, Institute of Neuroscience and Physiology, Sahlgrenska Academy at University of Gothenburg, Mölndal, Sweden; Clinical Neurochemistry Laboratory, Sahlgrenska University Hospital, Mölndal, Sweden; Department of Medical Statistics, London School of Hygiene & Tropical Medicine, London, United Kingdom; Department of Neurodegenerative Disease, UCL Queen Square Institute of Neurology, London, UK; Neurogenetics Laboratory, National Hospital for Neurology and Neurosurgery, University College London Hospitals NHS Foundation Trust, London WC1N 3BG, UK; National Prion Clinic, National Hospital for Neurology and Neurosurgery, University College London Hospitals NHS Foundation Trust, London WC1N 3BG, UK; MRC Prion Unit at UCL, UCL Institute of Prion Diseases, 33 Cleveland Street, London W1W 7FF, UK; VIB-KU Leuven Center for Brain & Disease Research, Leuven, Belgium; Department of Neurosciences, Leuven Research Institute for Neuroscience and Disease (LIND), KU Leuven, Leuven, Belgium

## Abstract

*In-vitro* studies of autosomal dominant Alzheimer’s disease (ADAD) implicate longer Aβ peptides in pathogenesis, however less is known about the behaviour of ADAD mutations *in-vivo*. In this cross-sectional cohort study, we used liquid chromatography-tandem mass spectrometry to analyse 66 plasma samples from ADAD family members who were at-risk of inheriting a mutation or were already symptomatic. We tested for differences in plasma Aβ42:38, 38:40 and 42:40 ratios between *Presenilin1 (PSEN1)* and *Amyloid Precursor Protein (APP)* carriers. We examined the relationship between plasma and *in-vitro* models of Aβ processing and, among *PSEN1* carriers, tested for associations with parental age at onset (AAO). 39 participants were mutation carriers (28 *PSEN1* and 11 *APP).* Age- and sex-adjusted models showed marked differences in plasma Aβ between *APP* and *PSEN1*: higher Aβ42:38 in *PSEN1* versus *APP* (p<0.001) and non-carriers (p<0.001); higher Aβ38:40 in *APP* versus *PSEN1* (p<0.001) and non-carriers (p<0.001), while Aβ42:40 was higher in *APP* and *PSEN1* compared to non-carriers (both p<0.001). Aβ profiles were reasonably consistent in plasma and cell lines. Within *PSEN1*, sex-adjusted models demonstrated negative associations between (i)Aβ42:40 (ii)Aβ42:38 and parental AAO. *In-vivo* differences in Aβ processing between *APP* and *PSEN1* provide insights into ADAD pathophysiology which can inform therapy development.

## INTRODUCTION

Understanding Alzheimer’s disease (AD) pathogenesis is critical to realising disease-modifying treatments. Autosomal dominant Alzheimer’s disease (ADAD), caused by mutations in presenilin 1/2 *(PSEN1/2)* or amyloid precursor protein (*APP)*, is a valuable model for characterising the molecular drivers of AD and its clinical heterogeneity (Ryan *et al.*, 2016).

PSEN1, the catalytic subunit of γ-secretase, sequentially cuts APP: the first endopeptidase cleavage generates an intracellular domain and a long membrane-associated amyloid-beta (Aβ) peptide, either Aβ49 (major product) or Aβ48 (minor product) (Sato *et al.*, 2003). Subsequent proteolytic cleavage of Aβ largely occurs down two alternative pathways: Aβ49>46>43>40 and Aβ48>45>42>38 (Takami *et al.*, 2009). As Aβ49 is the predominant endopeptidase cleavage product, normal processing of APP largely leads to Aβ40 formation (Sato *et al.*, 2003). Pathogenic ADAD mutations alter APP processing resulting in more, and/or longer, aggregation prone, Aβ peptides which accelerate cerebral amyloid accumulation leading to typical symptom onset in 30s to 50s (Bateman *et al.*, 2012; Chávez-Gutiérrez *et al.*, 2012).

Both *APP* and *PSEN1/2* mutations increase production of longer (e.g. Aβ42) relative to shorter (e.g. Aβ40) peptides (Chávez-Gutiérrez *et al.*, 2012). However, there are intriguing inter-mutation differences in Aβ profiles. *PSEN1* mutant cell lines produce increased Aβ42:38 ratios reflecting impaired γ-secretase processivity (Chávez-Gutiérrez *et al.*, 2012; Arber *et al.*, 2019). In contrast, *APP* mutations, located around the γ-secretase cleavage site, increase the Aβ38:40 ratio; consistent with preferential processing down the Aβ48 pathway (Chávez-Gutiérrez *et al.*, 2012; Arber *et al.*, 2019). To date, studies examining the influence of ADAD genotypes on Aβ ratios *in-vivo* have been lacking.

Increasingly sensitive mass spectrometry-based assays now make it possible to measure concentrations of different Aβ moieties in plasma (Schindler *et al.*, 2019). Therefore, we aimed to analyse plasma Aβ levels in an *APP* and *PSEN1* cohort, explore influences of genotype and clinical stage and examine relationships between ratios and age at onset (AAO) and also assess consistency with *in-vitro* models of Aβ processing.

## METHODS

### Study design and participants

We recruited 66 participants from UCL’s longitudinal ADAD study; details described previously (Ryan *et al.*, 2016). Samples were collected from August 2012 to July 2019 and concomitantly a semi-structured health questionnaire and clinical dementia rating (CDR) scale were completed (Morris, 1993). Estimated years to/from symptom onset (EYO) were calculated by subtracting parental AAO from the participant’s age. Participants were defined as symptomatic if global CDR was >0 and consistent symptoms of cognitive decline were reported. ADAD mutation status, determined using Sanger sequencing, was provided only to statisticians, ensuring blinding of participants and clinicians. The study had local Research Ethics Committee approval; written informed consent was obtained from all participants or from a consultee if cognitive impairment precluded informed consent.

### Measurement of plasma Aβ levels

EDTA plasma samples were processed, aliquoted, and frozen at −80 °C according to standardised procedures and shipped frozen to the Clinical Neurochemistry Laboratory, Sahlgrenska University Hospital for analysis blinded to participants’ mutation status and diagnosis. Analysis of processed samples was performed using a liquid chromatography-tandem mass spectrometry (LC-MS/MS) method using an optimized protocol for immunoprecipitation for improved analytical sensitivity (Appendix 1) (Pannee *et al.*, 2014). Pooled plasma samples were used to track assay performance; intra- and inter-assay coefficients of variation were <5%.

### Correlation of Aβ ratios in plasma and in induced pluripotent stem cell (iPSC) neurons

A sub-study was conducted to compare Aβ profiles in plasma and in iPSC-derived neurons. Aβ ratio profiles were compared based on mutation for 8 iPSC mutant lines; mutations tested were APP V717I (n=2), PSEN1 Intron 4 (n=1), Y115H (n=1), M139V (n=1), R278I (n=1) and E280G (n=2). Ideally plasma and iPSC samples were donated by the same participant. In cases where matched plasma from the same donor was not available, a plasma sample from a carrier of the same mutation, and if possible a member of the same family, was selected. Ratios for Aβ42:40, Aβ38:40 and Aβ42:38 were normalised by taking the ratio of the median ratio in controls for each experimental setting (n=27 non-carrier controls for plasma, n=5 iPSC lines from controls who were not members of ADAD families).

iPSC-neuronal Aβ was quantified as previously reported (Arber *et al.*, 2019). Briefly, iPSCs were differentiated to cortical neurons for 100 days and then 48 hour-conditioned culture supernatant was centrifuged to remove cell debris and Aβ analysis was performed via electrochemiluminescence on the MSD V-Plex Aβ peptide panel (6E10), according to the manufacturer’s instructions.

### Statistical analysis

Summary descriptive statistics were calculated by mutation type (*PSEN1*, *APP,* non-carriers) and box plots produced for Aβ42:38, Aβ38:40 and Aβ42:40 ratios. These Aβ ratios are displayed on logarithmic scales since the choice of numerator and denominator in these ratios is arbitrary. Age- and sex-adjusted differences were estimated between mutation type for each ratio; as were differences by clinical stage (presymptomatic vs symptomatic vs non-carriers) for *APP* in Aβ38:40 and Aβ42:40, and *PSEN1* in Aβ42:38 and Aβ42:40. These comparisons were made using mixed models including random intercepts for clusters comprising individuals from the same family and group, with random intercept and residual variances allowed to differ for the groups being compared. Pairwise comparisons were only carried out if a joint test provided evidence of differences. Ratios were log-transformed; estimated coefficients were back-transformed to multiplicative effects.

In *PSEN1* participants, mixed models with a random effect for family investigated sex-adjusted relationships of Aβ42:38 and Aβ42:40 with parental AAO; for the former model there was evidence to also include a quadratic term for parental AAO. In each analysis the estimated geometric mean ratio (and 95% confidence interval) was plotted against parental AAO standardising to the gender mix of the sample (54% female/46% male) (Fig. 2C, 2D). Additionally, the models were rerun adjusting separately for EYO and age. Associations between ratio levels and parental AAO were not tested for in *APP* carriers due to small sample size and limited array of mutations (Supplementary Table 1).

Spearman correlation coefficients were calculated to assess the association between plasma and iPSC-neuron Aβ ratios.

Analyses were performed using Stata v16.

### Data availability

Data will be made available upon reasonable request to qualified investigators, adhering to ethical guidelines.

## RESULTS

Demographic and clinical characteristics are presented in Table 1: 39 mutation carriers (28 *PSEN1*, 11 *APP*); 27 non-carriers. *APP* mutations investigated were located within the carboxyterminal region of Aβ, the site of the γ-secretase cleavage (Supplementary Table 1).

Age- and sex-adjusted models showed marked differences in plasma Aβ between *APP* and *PSEN1* mutation carriers. The geometric mean of the Aβ42:38 ratio was higher in *PSEN1* compared to both *APP* carriers (69% higher, 95%CI 39%, 106%; p<0.001) and to non-carriers (64% higher, 95%CI 37%, 98%; p<0.001), while there was no evidence of a difference between *APP* carriers and non-carriers (p= 0.60) (Fig. 1A). The geometric mean of Aβ38:40 ratio was twice as high (101% higher) in *APP* carriers compared to *PSEN1* carriers (95%CI 72%, 135 %; p<0.001) and 61% higher in *APP* carriers compared to non-carriers (95%CI 41%, 84%; p<0.001), while in *PSEN1* carriers the geometric mean of the Aβ38:40 ratio was 20% lower than in non-carriers (95%CI 10%, 29%, p<0.001) (Fig. 1B). Plasma Aβ42:40 ratios were raised in both *APP* and *PSEN1*; compared to non-carriers the geometric mean was estimated to be 61% higher (95%CI 44%, 80%; p<0.001) in *APP* and 31% higher (95%CI 16%, 50%; p<0.001) in *PSEN1* (Fig. 1C). There were also significant differences in the Aβ42:40 ratio between the *APP* and *PSEN1* groups; the estimated geometric mean was 22% higher (95%CI 8%, 38%; p=0.001) in *APP* carriers compared to *PSEN1* carriers.

**Figure 1:**
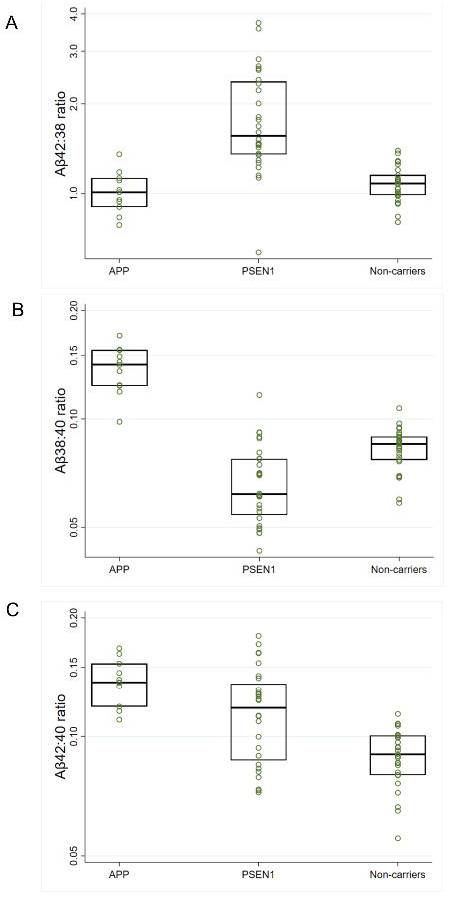
Box plots for observed plasma Aβ ratios across the three groups. Mutation carriers were divided into *APP* and *PSEN1* carriers. The unadjusted plasma (**1A**) Aβ42:38, (**1B)** Aβ38:40 and **(1C)** Aβ42:40 ratios are shown with the y-axis on a logarithmic scale. Boxes show the median and first and third quartiles. Dots represent individual observations.

Within *PSEN1,* the geometric mean Aβ42:40 ratio was 24% higher (95%CI 2%, 52%; p=0.03) in symptomatic compared to presymptomatic carriers; the Aβ42:38 ratio was also (non-significantly) 27% higher (95%CI 6% lower, 70% higher; p=0.11) in symptomatic compared to presymptomatic *PSEN1* carriers (Supplementary Fig. 1). No significant differences were observed in either the Aβ42:40 or the Aβ38:40 ratio between presymptomatic and symptomatic *APP* carriers (p>0.52).

Across *PSEN1* carriers, sex-adjusted models demonstrated negative associations between both the Aβ42:40 (p=0.003) and Aβ42:38 ratios (p<0.001) and parental AAO, higher ratios being associated with younger age at parental onset (Fig. 2). These associations remained statistically significant when additionally adjusting for EYO (p=0.01, p<0.001, respectively) or age (p=0.005, p<0.001, respectively). For Aβ42:38 there was evidence to include a quadratic term for parental AAO (p=0.004), resulting in the negative association with geometric mean of Aβ42:38 being most pronounced at younger parental AAO and reducing as parental AAO increased (Fig. 2C). In contrast, we did not find evidence to support the use of a quadratic term for Aβ42:40; a one-year increase in parental AAO was associated with a 1.8% reduction (95% CI: 0.6%, 3.0%) in the Aβ42:40 ratio (Fig. 2D).

**Figure 2:**
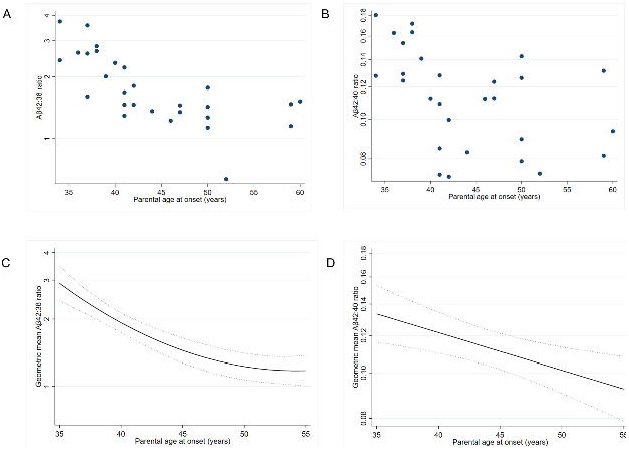
Plasma Aβ ratios against parental AAO in *PSEN1* carriers. Scatter plots of **(A)** observed plasma Aβ42:38 and **(B)** observed plasma Aβ42:40 values versus parental age at onset (AAO). Both scatter plots show values for *PSEN1* carriers only. Modelled geometric mean of **(C)** plasma Aβ42:38 and **(D)** plasma Aβ42:40 against parental AAO for a *PSEN1* carrier at EYO=0 (i.e. at expected symptom onset) standardised to the gender mix of the sample (54% female /46% male). Both models demonstrated significant negative associations – i.e. an earlier parental AAO was associated with a higher ratio. For Aβ42:40 the relationship between parental AAO and the plasma ratio was estimated to be constant across the age range shown: a one-year increase in parental AAO was associated with a 1.8% (95% CI: 0.6%, 3.0%) lower Aβ42:40 ratio. For Aβ42:38 the inclusion of a quadratic term resulted in the rate of change in the geometric mean reducing as parental AAO increased, for example: a 9.0% decrease (95% CI: 5.3%, 12.6%; p<0.001) for a one-year increase in parental AAO at parental AAO of 35 years; a 4.4% decrease (95% CI: 3.2%, 5.7%; p<0.001) at parental AAO of 45 years. The y-axis scale is logarithmic in all panes.

Aβ ratios in plasma and iPSC conditioned media were highly associated for both Aβ42:40 (rho=0.86, p=0.01) and Aβ38:40 (rho=0.79, p=.02), somewhat less so for Aβ42:38 (rho=0.61, p=0.10 (Fig. 3). While we did not observe perfect agreement in the Aβ42:38 ratio between plasma and iPSC lines (shown by solid line, Fig. 3), the direction of change in this ratio, i.e. either increased or decreased when compared to controls, was largely consistent across media.

**Figure 3:**
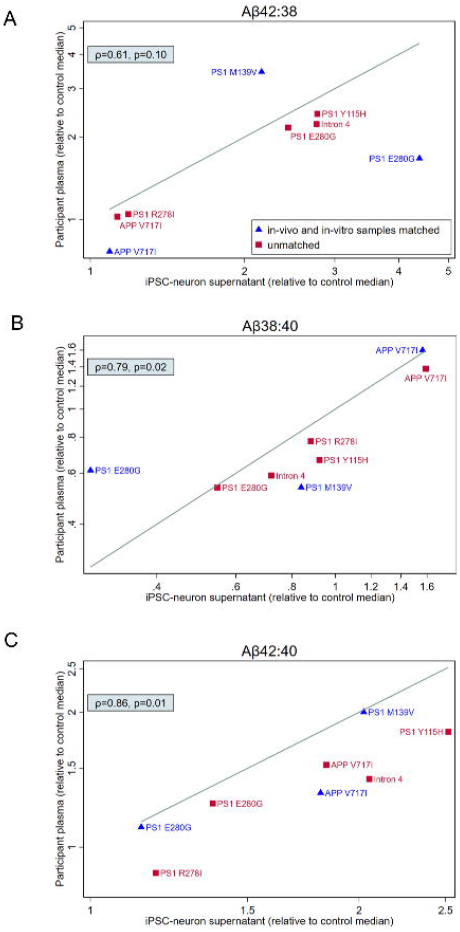
Comparison of Aβ processing *in-vivo* and *in-vitro.* Scatterplots comparing Aβ ratios profiles in plasma and iPSC derived neurons for eight mutation carriers. One to one comparison of Aβ ratios normalised to the median of controls for each experimental setting (n=27 non-carrier controls for plasma, n=5 iPSC lines from controls who were not members of ADAD families); values >1 indicate higher ratio in mutation carrier compared to median of controls whereas values <1 indicate lower ratio in mutation carrier compared to median of controls. The line displayed on each scatterplot represents line of perfect agreement i.e. x=y. Spearman rho and the associated p-value are shown for each scatter plot. Matched samples (plasma and iPSC samples donated by the same donor) are identified with triangle symbols. Unmatched samples (plasma and iPSC samples donated by different participants who carry the same mutation, and where possible are members of the same family) are identified by square symbols. The y-axis scale is logarithmic in all panes.

## DISCUSSION

In this study we found elevations in the plasma Aβ42:40 ratio in both *APP* and *PSEN1* carriers compared to non-carriers and differences in Aβ ratios across genotypes: Aβ42:38 ratios were higher in *PSEN1* vs. *APP*, Aβ38:40 ratios were higher in *APP* vs. *PSEN1*. Importantly, more aggressive *PSEN1* mutations (those with an earlier age at onset) had higher Aβ42:40 and Aβ42:38 ratios – *in-vivo* evidence of the pathogenicity of these peptide ratios.

These results offer insights into the pathobiology of ADAD and differential effects of *APP/PSEN1* genotype. Increased Aβ42:38 in *PSEN1* may be attributed to reduced conversion of Aβ42 (substrate) to 38 (product) relative to non-carriers – in contrast *APP* carriers showed near identical Aβ42:38 ratios compared to non-carriers. Additionally, we found an association with the relative increase of Aβ42 compared to shorter moieties (Aβ38, Aβ40) and the estimated timing of disease onset within *PSEN1* carriers. Importantly associations between Aβ42:38 and Aβ42:40 and parental AAO remained significant when adjusting for EYO; increasing the likelihood that these ratios represent key molecular drivers of disease onset as opposed to the confounding effects of disease stage. Our *in-vivo* results recapitulate cell-based findings of reduced efficiency of γ-secretase processivity in *PSEN1* mutations (Szaruga *et al.*, 2015, 2017*b*; Arber *et al.*, 2019). This inefficiency is attributed to impaired enzyme-substrate stability causing premature release of longer, aggregation-prone Aβ peptides (Chávez-Gutiérrez *et al.*, 2012; De Strooper and Karran, 2016).

The estimated timing of symptom onset serves as an indicator of disease severity, with a younger parental AAO suggestive of a more deleterious *PSEN1* mutation. When modelling plasma Aβ42:38, a read-out of the efficiency of the fourth γ-secretase cleavage cycle, we found deceleration in the rate of change as parental AAO increases. This finding further supports the central pathogenic role of γ-secretase processivity in ADAD, especially in younger onset, aggressive forms of *PSEN1.*

In *APP* carriers, production of Aβ38 relative to Aβ40 was increased compared to non-carriers and *PSEN1* carriers. This is consistent with a shift in the site of endopeptidase-cleavage causing increased generation of Aβ48; the precursor substrate in the Aβ38 production line. Our study included *APP* mutations (Supplementary Table 1) located around the γ-secretase cleavage site. Previous cell-based work involving mutations around this site also found evidence of increased trafficking of APP down the Aβ48 pathway (Chávez-Gutiérrez *et al.*, 2012, Szaruga *et al.*, 2017*b*; Arber *et al.*, 2019). In contrast, *APP* duplications or mutations near the beta-secretase site are associated with non-differential increases in Aβ production (Hunter and Brayne, 2018).

Heterogeneity is also seen within *PSEN1*: mutations have differential effects on γ-secretase processivity and occasionally also on the proportion of substrate being trafficked down Aβ pathways (Fernandez *et al.*, 2014; Arber *et al.*, 2019). Additionally, studies of γ-secretase processivity demonstrate associations between substrate length and enzyme-substrate stability (Szaruga *et al.*, 2017*b*). Pathogenic *PSEN1* mutations further destabilise, albeit to differing extents, the enzyme-substrate complex; increasing the likelihood of release of longer (>Aβ43 peptides) (Szaruga *et al.*, 2017*b*). Therefore, declines in Aβ38:40 in *PSEN1* relative to non-carriers may be due to mutation effects on catalysis within processing pathways and/or across cleavage sites.

Our results support the hypothesis that *APP* and *PSEN1* mutations increase *in-vivo* production of long Aβ peptides (Aβ ≥42) relative to Aβ40. This is consistent with cell- and blood-based studies in ADAD (Reiman *et al.*, 2012; Fagan *et al.*, 2014). The increase in the Aβ42:40 ratio in plasma contrasts with the well-established decline in this ratio in CSF in ADAD; reductions in CSF levels are attributed to sequestration or “trapping” of longer (more aggregation-prone) peptides within cerebral amyloid plaques (Fagan *et al.*, 2014; Fortea *et al.*, 2020). Thus, plasma Aβ, which reflects both central and peripheral production, appears to be less susceptible to the confounding effect of sequestration than CSF Aβ. In addition, we show consistency in Aβ ratio profiles between plasma and cell media, although it is important to avoid over interpreting this finding in such small numbers. Taken together, these findings suggest that plasma Aβ ratios may provide ideal, easily accessible biomarkers of Aβ processing as well as *PSEN1/APP* gene function. Given ongoing efforts to halt AD progression by targeting amyloid, such measures are valuable, and may become even more so as we enter an era of personalised medicine and gene-based therapies.

We show some evidence of change across disease stage with increased Aβ42:40 in symptomatic individuals compared to presymptomatic *PSEN1* carriers. The reason for this is unclear and should be treated cautiously, especially given the small numbers, however it is interesting to note that increases in plasma Aβ42 after symptom onset in Down syndrome have been reported (Fortea *et al.*, 2020). Heterogeneity in pathobiology may contribute to differences across disease stage; intra-mutation fluctuations in Aβ ratios have previously been associated with variability in *PSEN1* maturation (Arber *et al.*, 2019). In addition, the pathogenic consequences of ADAD may contribute to the changes seen, with recent evidence suggesting γ-secretase processivity declines with advancing Braak stage (Kakuda *et al.*, 2020).

Our study has limitations including that the sample size was small due to the relative rarity of ADAD, however this cohort does contain a reasonably wide array of *PSEN1* and *APP* mutations. We did not have paired CSF (important for clarifying relative contributions of central and peripheral Aβ production/clearance to plasma ratios), however we did compare Aβ ratios between plasma and iPSC conditioned media; reassuringly *in-vivo* and *in-vitro* results were reasonably consistent. In addition, we used parental age at symptom onset to estimate the timing of disease onset. While parental AAO is a reasonable estimate, it is not without error due both to variability in age at onset between family members and to imprecision in determining the exact timing of symptom onset in a preceding, now often deceased, generation (Ryman *et al.*, 2014; Ryan *et al.*, 2016). Finally, future studies should measure Aβ moieties longer than Aβ42 to better clarify AD pathogenesis. Nonetheless, these findings offer insights into ADAD pathophysiology *in-vivo* that may inform therapy development.

## Contributors

AOC, NCF, HZ developed the study concept. AOC, PSJW, NSR, and HR contributed to recruitment. Data were collected by AOC, PSJW, HR, CA, NW and NSR. Blood samples were processed by AJH, EA, and IS. The immunoprecipitation mass spectrometry method was developed by JP, KB, HZ, and EP. JP analysed the plasma samples. TP, JMN and CF carried out the statistical analysis. SM and JMP contributed to the genetic analysis. TP and CF created the figures. AOC, JP, TP, NSR, CA, CF, SW, LCG, KB, HZ, and NCF interpreted the data. AOC and NCF drafted the initial manuscript. All authors reviewed and edited the manuscript and critically revised it for intellectual content.

## Acknowledgements

AOC is supported by an Alzheimer’s Society clinical research training fellowship (AS-CTF-18-001). CA is supported by a fellowship from the Alzheimer’s Society (AS-JF-18-008) and SW is supported by an Alzheimer’s Research UK Senior Research Fellowship (ARUK-SRF2016B-2). IS is supported by the UK Dementia Research Institute which receives its funding from DRI Ltd, funded by the UK Medical Research Council, Alzheimer’s Society and Alzheimer’s Research UK. PSJW is supported by an MRC Clinical Research Training Fellowship. NSR is supported by a University of London Chadburn Academic Clinical Lectureship. HZ is a Wallenberg Scholar supported by grants from the Swedish Research Council (#2018-02532), the European Research Council (#681712), Swedish State Support for Clinical Research (#ALFGBG-720931) and the UK Dementia Research Institute at UCL. KB is supported by the Swedish Research Council (#2017-00915), the Alzheimer Drug Discovery Foundation (ADDF), USA (#RDAPB-201809-2016615), the Swedish Alzheimer Foundation (#AF-742881), Hjärnfonden, Sweden (#FO2017-0243), and European Union Joint Program for Neurodegenerative Disorders (JPND2019-466-236). CF, JMN and TP’s academic collaboration with the Dementia Research Centre, UCL is supported by a grant to the DRC from Alzheimer’s Research UK. NCF acknowledges support from Alzheimer’s Research UK, the UK Dementia Research Institute and the NIHR UCLH Biomedical Research Centre. This work was supported by the NIHR UCLH/UCL Biomedical Research Centre, the Rosetrees Trust, the MRC Dementia Platform UK and the UK Dementia Research Institute at UCL which receives its funding from UK DRI Ltd, funded by the UK Medical Research Council, Alzheimer’s Society and Alzheimer’s Research UK, and the Swedish state under the agreement between the Swedish government and the County Councils, the ALF-agreement (#ALFGBG-715986). Professor Nick Fox had full access to all the data in the study and takes responsibility for the integrity of the data and the accuracy of the data analysis.

