## Supplementary material for "Plasma Aβ ratios in autosomal dominant Alzheimer’s disease: the influence of genotype"

| **Gene** | **Mutation** | **Number of individuals** |
| --- | --- | --- |
| APP | p.Thr719Asn | 1 AR |
|  | p.Val717Gly | 2 S |
|  | p.Val717Ile | 1 S, 7 AR |
|  | p.Val717Leu | 2 S, 2 AR |
| PS1 | Intron 4 | 2 S, 5 AR |
|  | p.Ala79Val | 1 S |
|  | p.Tyr115His | 1 S, 1 AR |
|  | p.Glu120Lys | 1 AR |
|  | p.Ser132Ala | 2 AR |
|  | p.Met139Val | 1 S, 1 AR |
|  | p.Val142Ile | 1 S |
|  | p.Met146Ile | 2 AR |
|  | p.Glu184Asp | 2 S, 4 AR |
|  | p.Ile202Phe | 4 AR |
|  | p.Gly206Ala | 1 S |
|  | p.His214Tyr | 3 AR |
|  | p.Ala246Glu | 2 AR |
|  | p.Pro264Leu | 2 AR |
|  | p.Pro267Ser | 1 S |
|  | p.Arg269His | 1 AR |
|  | p.Arg278Ile | 3 AR, |
|  | p.Glu280Gly | 2 S, 6 AR, |
|  | ΔE9* | 1 S |

**Table 1: The number of individuals from families with each mutation is given, divided in to symptomatic (S) or asymptomatic but at risk (AR).** Details relating to how many at risk participants for each mutation were mutation carriers is not given to ensure it is not possible the mutation status of any at risk individual to be revealed/deduced. ** The exon 9 deletion (NM_000021.3:c.869-1G>T; p.Ser290Cys;Thr291_Ser319del) commonly referred to as ΔE9.

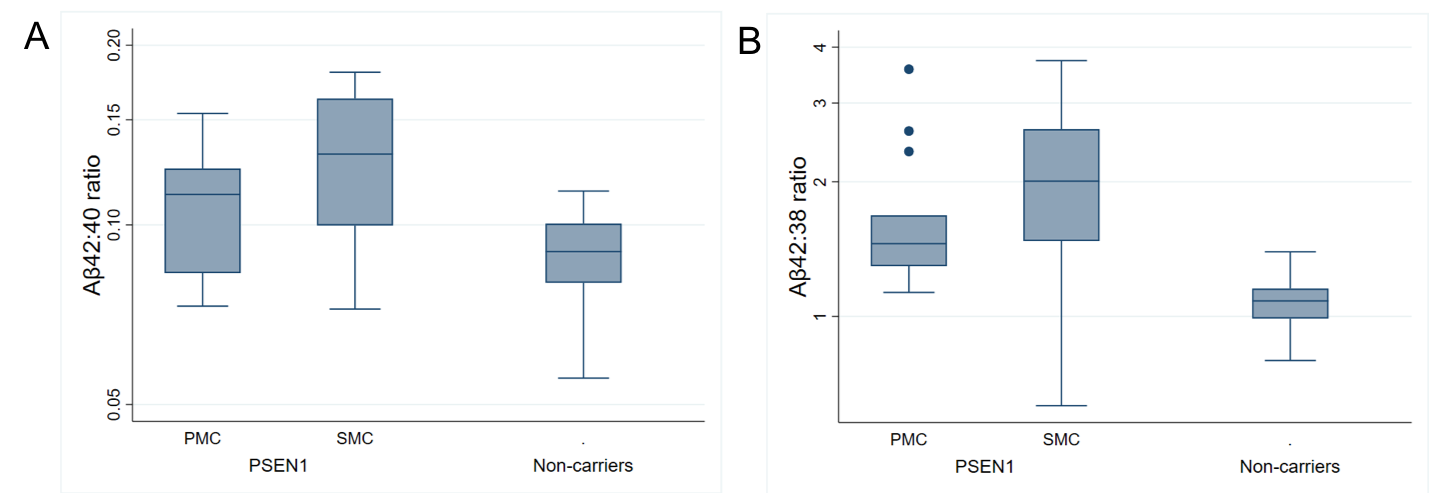

**Figure 1:** **Box and whisker plots for observed baseline plasma Aβ ratios**. *PSEN1* mutation carriers were divided into presymptomatic (PMC) and symptomatic (SMC) groups. Boxes show the median and first and third quartiles; whiskers show data within 1·5 IQR of the median; dots represent outliers. The y-axis scale is logarithmic. All group comparisons were carried out on log-transformed ratios, adjusting for age and sex. For each analysis a joint test provided evidence of a difference (p<0.001).

**Appendix 1:**

**Measurement of plasma Aβ levels**

Calibrators were prepared using recombinant Aβ1-38, Aβ1-40 and Aβ1-42 (rPeptide) added to 8% bovine serum albumin in phosphate-buffered saline. Recombinant 15N uniformly labelled Aβ1-38, Aβ1-40 and Aβ1-42 (rPeptide) were used as internal standards (IS), added to samples and calibrators prior to sample preparation. Aβ peptides were extracted from 250 µL human plasma using immunoprecipitation with anti-β-Amyloid 17-24 (4G8) and anti-β-Amyloid 1-16 antibodies (6E10, both BioLegend®) coupled to Dynabeads™ M-280 Sheep Anti-Mouse IgG magnetic beads (Thermo Fisher Scientific). Immunoprecipitation was performed using a KingFisher™ Flex Purification System (Thermo Fisher Scientific). Analysis of processed samples was performed using liquid chromatography-tandem mass spectrometry (LC-MS/MS) on a Dionex Ultimate LC-system and a Thermo Scientific Q Exactive quadrupole-Orbitrap hybrid mass spectrometer. Chromatographic separation was achieved using basic mobile phases and a reversed-phase monolith column at a flow rate of 0.3 mL/min. The mass spectrometer operated in parallel reaction monitoring (PRM) mode was set to isolate the 4+ charge state precursors of the Aβ peptides. Product ions (14-15 depending on peptide) specific for each precursor was selected and summed to calculate the chromatographic areas for each peptide and its corresponding IS. The area ratio of the analyte to the internal standard in unknown samples and calibrators was used for quantification.
